# Repeatable quantum-hardware execution of a fast local-topology surrogate for hyperthermal sarcomeric oscillations

**DOI:** 10.64898/2026.04.02.716025

**Authors:** Seine A. Shintani

## Abstract

Cardiac contraction depends on sarcomeres acting locally, but a beating cardiomyocyte is not a perfectly uniform lattice: neighbouring sarcomeres can rapidly rephase while the cell keeps a slower rhythm. Hyperthermal sarcomeric oscillations (HSOs), a warmed-cardiomyocyte phenomenon previously identified at the sarcomere level, provide a compact experimental case of this mesoscale coordination problem. I recast the experimentally defined five-sarcomere HSO topology as a four-qubit quantum-hardware state space: four neighbouring-pair phase relations define 16 basis states, each state is evolved with a short nearest-neighbour two-step Trotter kernel, and IBM EstimatorV2 reads out physiology-linked diagonal observables. These readouts retain the HSO meanings of pattern persistence, one-link reconfiguration, anti-phase-rich occupancy, mismatch placement, and a topology-derived synchrony proxy (Stopo). The locked `control_base` lane acted as a reproducible hardware anchor. Across three independent 4096-shot candidate panels it produced weighted stay 0.8395 ± 0.0016, single-link fraction 0.9134 ± 0.0049, anti-phase-rich occupancy 0.4517 ± 0.0019, transition edge proxy 0.5437 ± 0.0016, and Stopo 0.2954±0.0002. Candidate panels then used the same lane as a model-family stress test. The overmixed reference marked a high-transition boundary, with transition mass 0.3837 ± 0.0313, single-link fraction 0.8248 ± 0.0198, and anti-phase-rich occupancy 0.4302 ± 0.0036. The edge-like representative `edge_candidate_alt3` occupied an intermediate hardware regime, with transition mass 0.2781±0.0038 and anti-phase-rich occupancy 0.4447±0.0020, while preserving statewise ordering for anti-phase-rich occupancy and Stopo. A fixed-prior closed-unitary bound placed the anti-phase-rich ceiling at 0.4642, below the biological HSO reference of 0.509, indicating model headroom rather than hardware readout failure. Thus the device functions as a hardware-aware physiological testbed: HSO-like coordination is constrained local reconfiguration, whereas excessive mixing erodes one-link and anti-phase-rich readouts.

**Significance statement:** This study starts from a concrete living-cell phenomenon rather than from an abstract small circuit. Hyperthermal sarcomeric oscillations are warmed-cardiomyocyte sarcomere dynamics in which a beat-scale rhythm persists while neighbouring sarcomeres rapidly redistribute phase. I map a five-sarcomere HSO segment to a four-qubit nearest-neighbour topology and read it out with physiology-linked observables rather than a generic circuit score. The central result is repeatable hardware execution of biologically meaningful local-state observables, and the physiological message is that HSO local grammar is constrained one-link-dominated rephasing, not maximal mixing or simple global synchrony.

## 1 Introduction

Cardiac contraction is generated by sarcomeres, but a beating cell is not a perfectly uniform lattice. Adjacent sarcomeres can differ in length, strain, phase, and mechanical history, and those differences feed into the amplitude and timing of the averaged cellular signal. Recent measurements make this point directly: stretch harmonizes sarcomere strain across cardiomyocytes, sarcomere-length variability persists within single cells, and in vivo nanoimaging links sarcomeric synchrony and asynchrony to ventricular pump performance (Li *et al*., 2023; Lookin *et al*., 2023; Kobirumaki-Shimozawa *et al*., 2021, 2024). Local nonuniformity is therefore a physiological variable, not a nuisance term to be averaged away.

A useful example of this local variable is hyperthermal sarcomeric oscillation (HSO), a warmed-cardiomyocyte phenomenon first identified as high-frequency sarcomeric auto-oscillation induced by heating in living neonatal cardiomyocytes (Shintani, 2015). In later HSO studies, the cell-level contraction rhythm remained homeostatic while the internal sarcomere signal showed rapid oscillation and amplitude modulation (Shintani *et al*., 2020; Shintani, 2022). This definition matters for the present work: HSO is not a generic “heated cell” label, but a sarcomere-level state in which beat-scale organization and fast local rephasing coexist. The warmed-cell context is mechanistically relevant because local heat pulses can trigger cardiomyocyte contraction without detectable calcium transients, microscopic heating activates cardiac thin filaments, and thermal activation shifts thin-filament regulatory equilibria toward activation (Oyama *et al*., 2012; Ishii *et al*., 2019, 2020). Cardiac activation also depends on cooperative thin-filament and thick-filament regulation and on length- and strain-dependent feedback (Solís and Solaro, 2021; Park-Holohan *et al*., 2021; de Tombe *et al*., 2010; Campbell, 2011). HSOs therefore place local sarcomere coordination in a physiologically interpretable regime: the system remains rhythmically organized, yet the internal contractile elements have room to redistribute phase.

Previous HSO studies established high-frequency sarcomeric auto-oscillation in living neonatal cardiomyocytes (Shintani, 2015), contraction-rhythm homeostasis during the HSO state (Shintani *et al*., 2020), amplitude modulation with chaotic properties while cycle timing remains stable (Shintani, 2022), and a periodic-chaotic or chaordic homeodynamic framing of the phenomenon (Shintani, 2025). More recent local-topology analyses reduced five consecutive sarcomeres to four neighbouring-pair phase relations and showed that HSO trajectories occupy a constrained 16-state topology dominated by Hamming-1 transitions and enriched anti-phase-rich occupancy (Shintani, 2026a). Related route analyses further suggested that mismatch placement along the five-sarcomere chain carries structured information (Shintani, 2026b). These findings define a compact mesoscale grammar: local HSO reconfiguration proceeds through small neighbour-to-neighbour changes rather than wholesale random resets.

This compact grammar creates a specific opportunity for quantum hardware. Many quantum-biology and quantum-biomedical proposals begin from molecules, sequences, or abstract networks (Emani *et al*., 2021; Outeiral *et al*., 2021; Cordier *et al*., 2022; Kösoglu-Kind *et al*., 2023; Roman-Vicharra and Cai, 2023; Li *et al*., 2024; Ghazi Vakili *et al*., 2025). The present study starts instead from an experimentally measured cell-physiology state space. Four neighbouring-pair relations naturally form four binary variables, and a nearest-neighbour quantum circuit naturally mirrors the local chain geometry. The aim is deliberately not to claim whole-cell simulation or quantum advantage. The scientific question is narrower and testable: can a biological local-state grammar be encoded, perturbed, and read out on real quantum hardware while preserving observables that retain physiological meaning?

I address that question by implementing a locked four-qubit HSO surrogate and a set of candidate kernels on IBM quantum hardware. The locked kernel tests repeatability and exact-vs-hardware ordering. The candidate panels interrogate the model family: a regular reference probes a conservative regime, an overmixed reference probes a high-transition boundary, and edge-like candidates test whether exact-screened intermediate regimes transfer to real hardware. Motivated by the general idea that useful dynamics can emerge between overly regular and overly mixed regimes (Goto *et al*., 2026), I use an operational criterion rather than a mechanistic claim: a useful HSO kernel should provide transition capacity while preserving one-link topology, anti-phase-rich readout, mismatch-placement information, and statewise interpretability on hardware. The resulting study turns local sarcomere coordination into an executable, hardware-testable physiological state space.

## 2 Results

### 2.1 A five-sarcomere segment is reduced to a four-qubit executable surrogate

The biological starting point is a five-sarcomere chain observed during HSO reanalysis. Each neighbouring pair was classified as in-phase or anti-phase, giving four binary relations: (1–2), (2–3), (3–4), and (4–5). These four relations form a 16-state local topology. I encoded the four relations into four qubits, with bit value 1 representing in-phase and 0 representing anti-phase (Figure 1). This is a relation-state encoding rather than a point-position encoding: a computational basis state is already a local HSO topology state.

**Figure 1.**
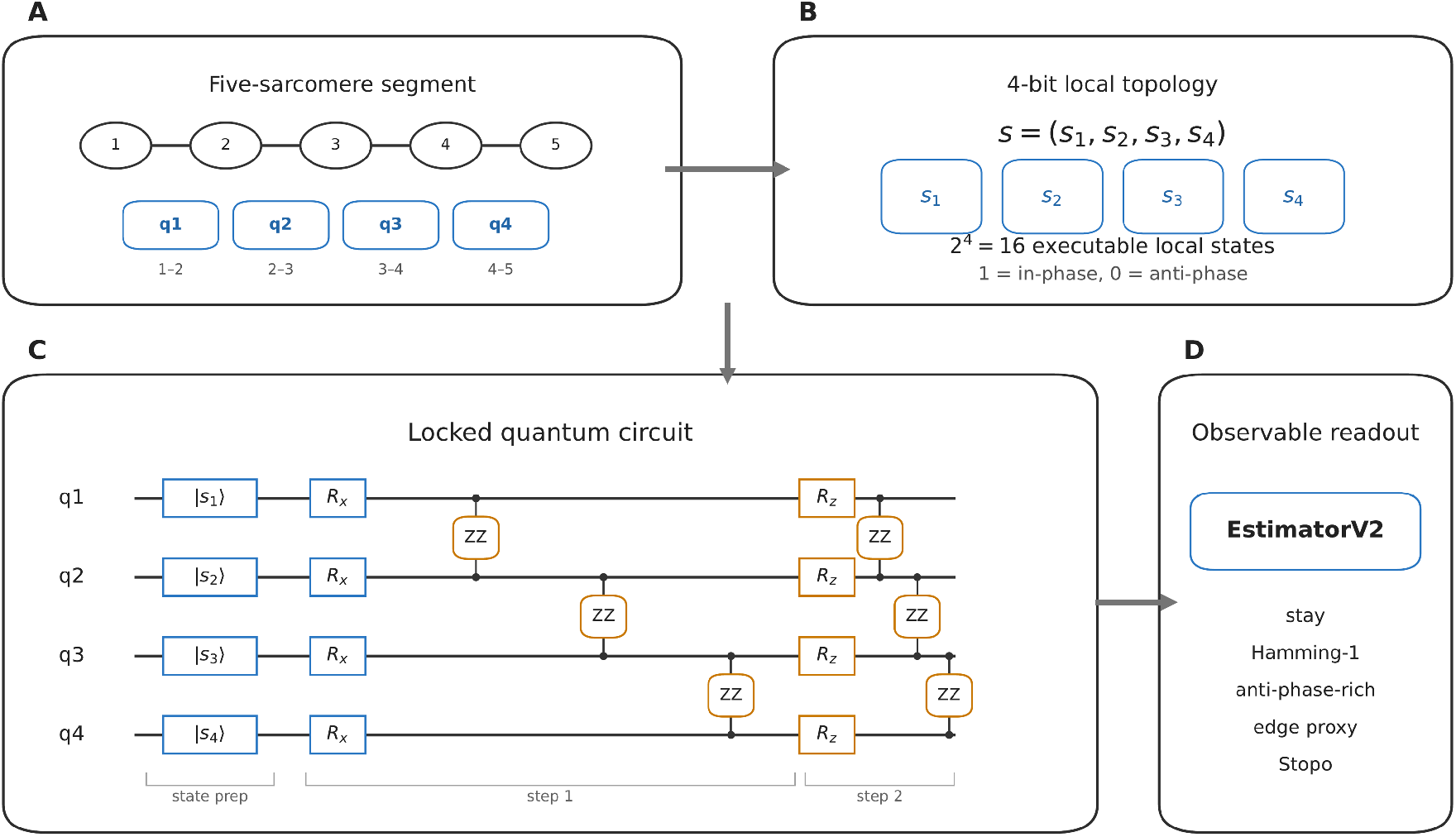
Mapping HSO local topology to a four-qubit executable surrogate. Five consecutive sarcomeres are reduced to four neighbouring-pair relations, and each relation is encoded as one qubit. The same 16 computational-basis states define the biological local-topology space and the quantum-circuit input space. Diagonal readout observables report stay, Hamming-distance transition structure, anti-phase-rich occupancy, mismatch placement, and Stopo.

For each of the 16 basis inputs, a short two-step Trotter circuit implemented a nearest-neighbour Hamiltonian kernel. Hardware readout used a diagonal observable pack designed around the HSO interpretation. Weighted stay measures persistence of the local topology. Hamming-1 mass measures one-link reconfiguration. Hamming *≥* 2 mass measures larger jumps. Anti-phase-rich occupancy reports whether at least three of the four neighbouring-pair relations are anti-phase. The edge proxy reports whether the minority mismatch pocket lies nearer the edge or the interior of the five-sarcomere segment. Stopo summarizes segment-mean synchrony after gauge-reconstructing the four pair relations into a five-sign pattern. This construction keeps the quantum-hardware output in the same language as the HSO local-topology analysis and avoids reducing the experiment to a generic fidelity score.

### 2.2 The locked control lane remains repeatable on IBM hardware

The locked lane, `control_base`, was designed as a stable hardware bridge. In the 512-shot repeatability panel, the same lane was run three times on `ibm_pittsburgh` using physical qubits [154, 153, 152, 151]. Across those three repeats, the hardware means were weighted stay 0.8506 ± 0.0041, single-link fraction 0.9075 ± 0.0094, anti-phase-rich occupancy 0.4519 ± 0.0022, transition edge proxy 0.5390 ± 0.0052, and Stopo 0.2946 ± 0.0008 (Figure 2).

**Figure 2.**
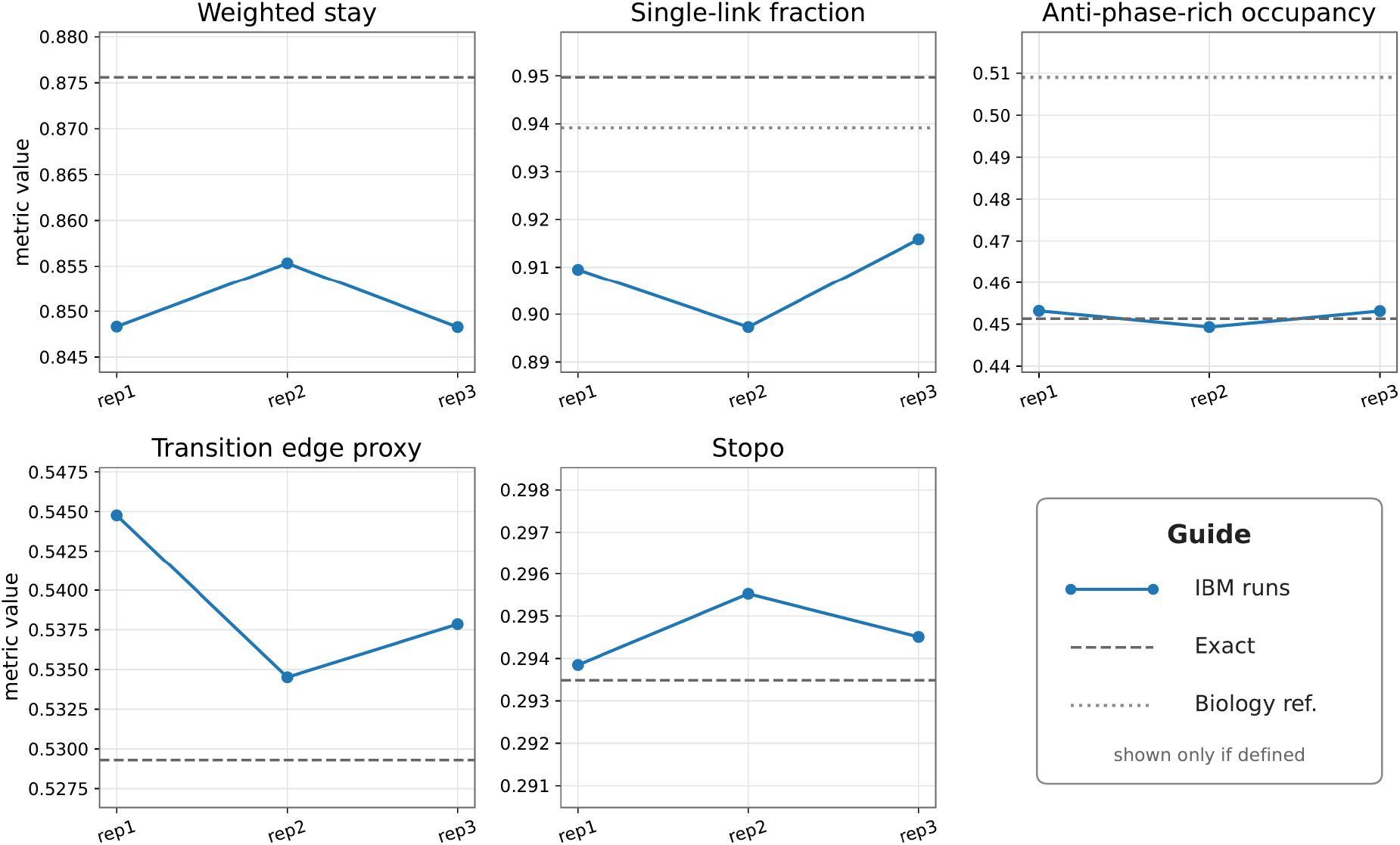
Repeat stability of the locked hardware lane. The same `control_base` lane was run three times on `ibm_pittsburgh` with 512 shots. Solid points and lines show IBM hardware repeats. The dashed line is the exact locked-surrogate reference. The dotted line marks the biology-side target where a target is defined from the earlier HSO reanalysis.

The 4096-shot candidate panels placed the same locked lane alongside candidate kernels. Across three independent 4096-shot panels containing `control_base`, the hardware means were weighted stay 0.8395±0.0016, single-link fraction 0.9134±0.0049, anti-phase-rich occupancy 0.4517±0.0019, transition edge proxy 0.5437 ± 0.0016, and Stopo 0.2954 ± 0.0002 (Table 2). The absolute stay level shifted modestly between the 512-shot and 4096-shot protocols, while the biology-linked readout pattern remained stable. Thus control_base functions as the current real-hardware anchor for this model family and provides the reference lane against which candidate kernels should be judged.

**Table 1.** Runtime environment and model metadata.

| Item | 512-shot repeatability panel | 4096-shot candidate panels |
| --- | --- | --- |
| Backend | `ibm_pittsburgh` | `ibm_pittsburgh` |
| Physical qubits | [154, 153, 152, 151] | [154, 153, 152, 151] |
| Primitive | EstimatorV2 | EstimatorV2 |
| Shots | 512 | 4096 |
| Resilience level | 1 | 1 |
| Optimization level | 1 | 1 |
| Trotter steps | 2 | 2 |
| Software environment | Qiskit 2.3.1; qiskit-ibm-runtime 0.45.1 | Qiskit 2.4.1; qiskit-ibm-runtime 0.47.0 |
| Role | Locked baseline repeatability | Hardware-aware candidate validation |

**Table 2.**
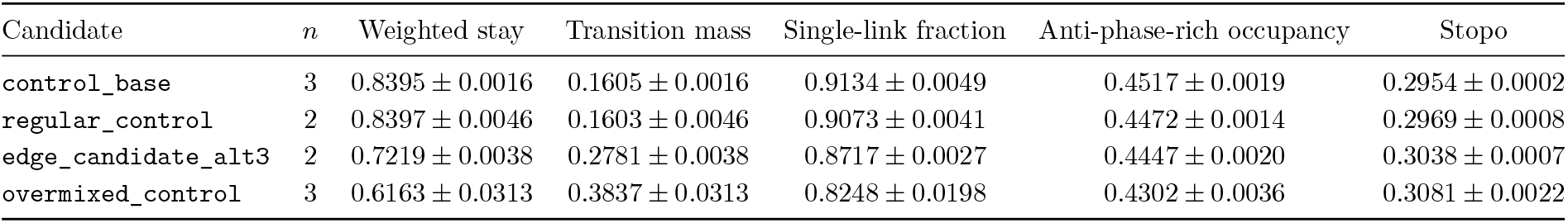
Repeated 4096-shot IBM candidate summaries. Values are mean ± sample SD across 4096-shot panels containing the candidate. *n* is the number of panels.

### 2.3 Statewise exact-vs-hardware structure is preserved in the locked execution

Aggregate means show repeatability, and statewise comparisons show that the hardware bridge preserves topology-dependent ordering across all 16 initial states. In the locked lane, Pearson correlation between exact and across-repeat hardware means was *r* = 0.945 for stay, 0.958 for Hamming-1 mass, 1.000 for anti-phase-rich occupancy, 0.941 for transition edge proxy, and 1.000 for Stopo. Mean absolute error was 0.0283, 0.0203, 0.0117, 0.0167, and 0.0070, respectively. These statistics ask whether each initial biological topology remains interpretable after hardware execution, not merely whether a single averaged number is close.

Figure 3 shows this agreement across the 16 initial states. State ordering is especially strong for anti-phase-rich occupancy and Stopo, while weighted stay and Hamming-1 mass carry the largest hardware shifts. The device therefore preserves the biologically relevant topology ordering while adding a modest transition bias characteristic of the real hardware execution.

**Figure 3.**
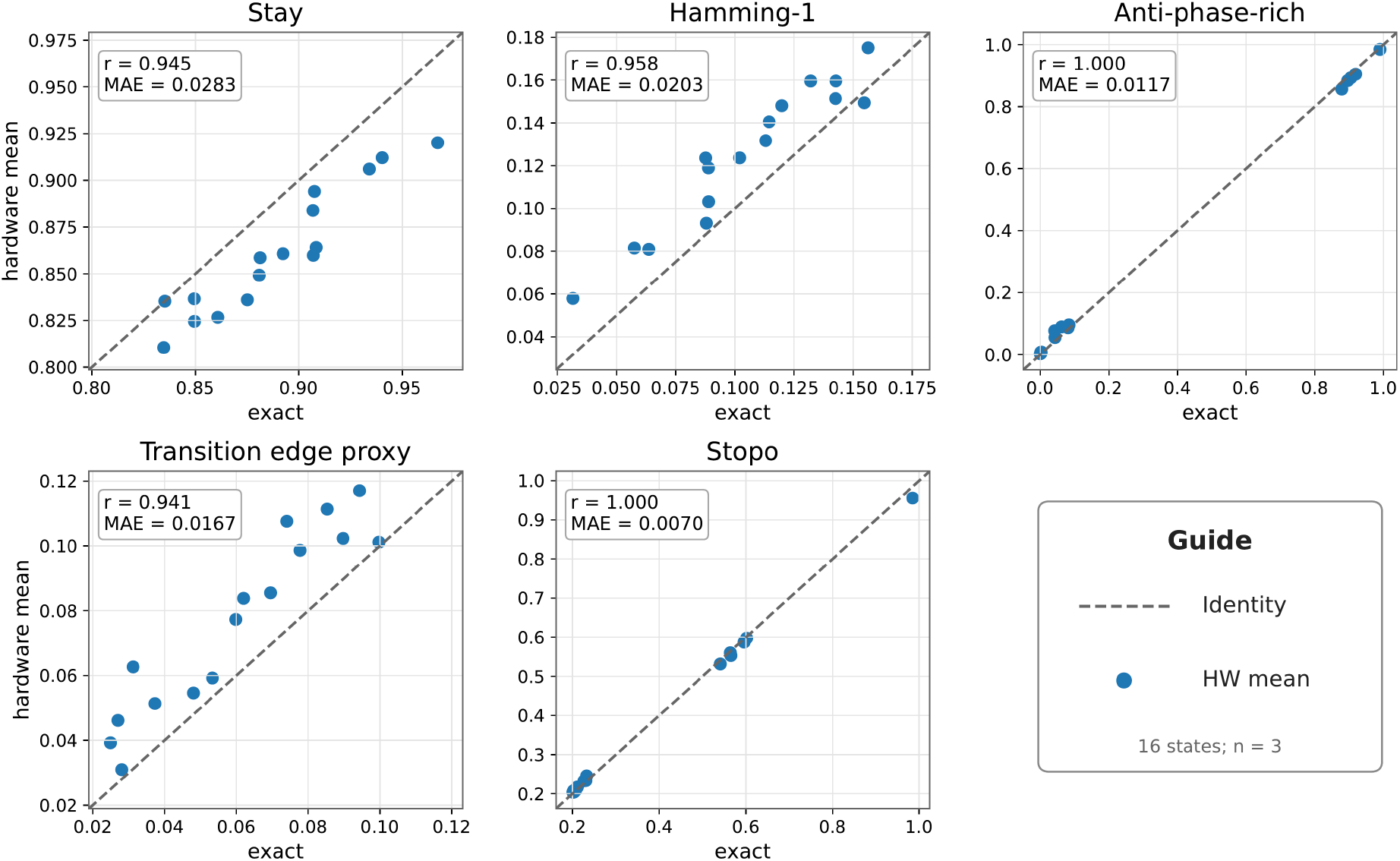
Statewise exact-vs-hardware agreement across the 16 initial topology states. Each point is the across-repeat hardware mean for one initial state plotted against the exact locked-surrogate value for the same state. The dashed line is the identity line. Agreement is strongest for anti-phase-rich occupancy and Stopo, and all five observables preserve clear statewise structure.

### 2.4 4096-shot candidate panels map conservative, edge-like, and high-transition regimes

The candidate-panel analysis introduced a compact model family around the locked baseline. The purpose was to span conservative, intermediate, and high-transition regimes with a small number of interpretable control knobs. The general form was

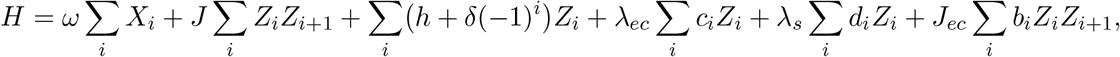

where *c* = (1, *−*1, *−*1, 1), *d* = (1, 1, *−*1, *−*1), and *b* = (1, *−*2, 1) represent simple edge-centre, side, and bond-asymmetry controls on the four-qubit line. These terms are low-dimensional control knobs for testing how local kernels move within the HSO observable space. They should not be read as molecular Hamiltonian terms; molecular or cellular interpretation is assigned to the resulting observable pattern rather than to an individual coefficient.

Exact screening classified candidate kernels as conservative, edge-like, or high-transition. The hardware panels then tested selected representatives on the same backend and physical layout with 4096 shots. The repeated summaries are shown in Figure 4 and Table 2. `regular_control` remained close to `control_base` in weighted stay and transition mass and preserved high single-link fraction, while anti-phase-rich occupancy sat below the baseline. `overmixed_control` defined the high-transition boundary: transition mass reached 0.3837 ± 0.0313, single-link fraction was 0.8248 ± 0.0198, and anti-phase-rich occupancy was 0.4302 ± 0.0036. This reference delineates the region in which transition capacity becomes strong enough to erode the one-link HSO topology.

**Figure 4.**
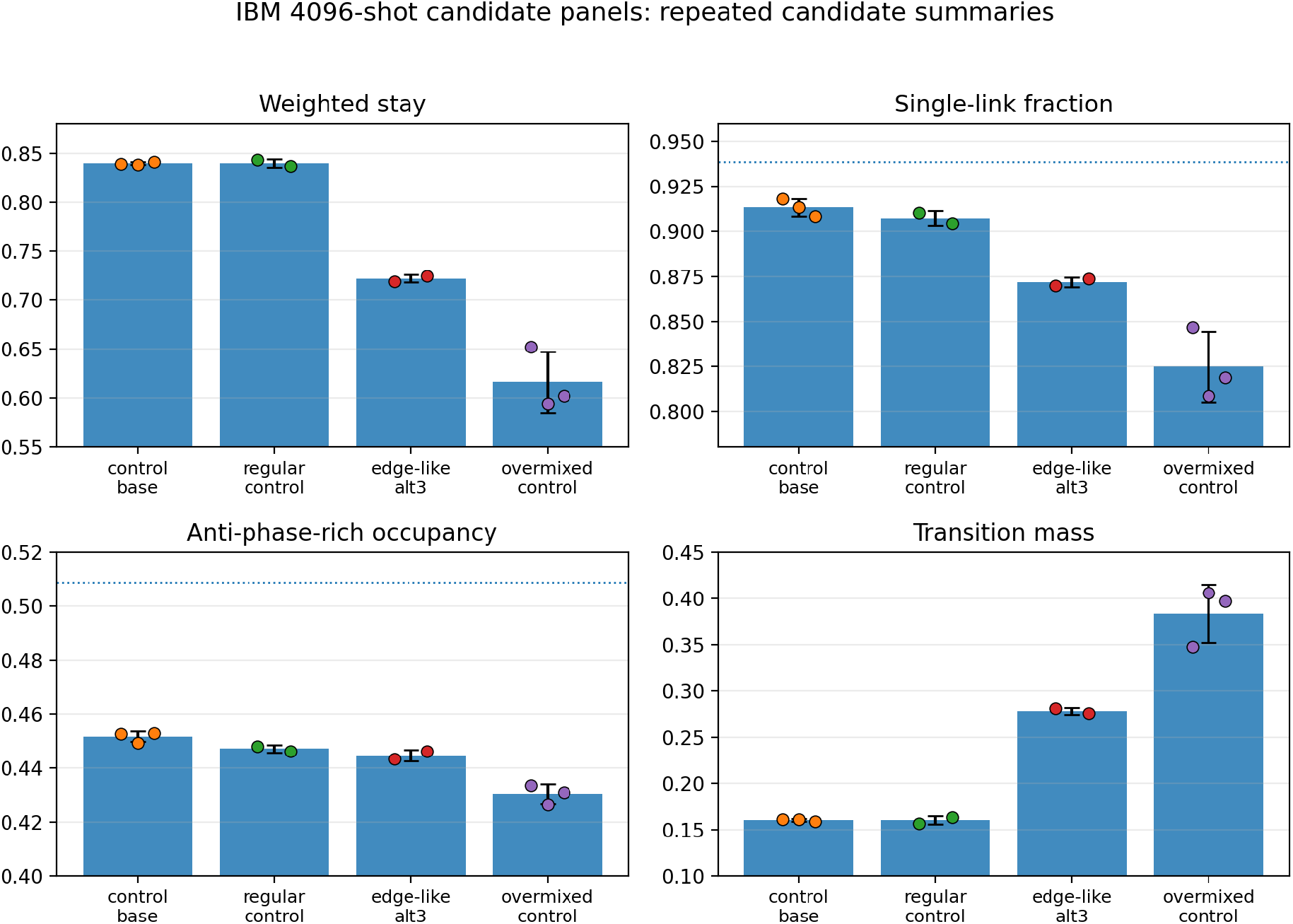
IBM 4096-shot candidate panels. Bars show mean ± sample SD for candidates appearing in repeated 4096-shot panels; overlaid points are individual IBM runs. `control_base` remains the strongest anchor. `regular_control` maps a conservative region. `edge_candidate_alt3` increases transition mass while preserving interpretable readouts. `overmixed_control` defines the high-transition boundary where single-link fraction and anti-phase-rich occupancy decline. Dotted guides mark biology-side reference values where defined.

The edge-like candidates provided the hardware-transfer information that turns exact screening into hardware-aware model selection. The initial exact-selected `edge_candidate` moved into a more transitional hardware regime. `edge_candidate_alt2` improved single-link retention relative to that first edge candidate. The most stable representative was `edge_candidate_alt3`. Across two 4096-shot runs, it gave weighted stay 0.7219 ± 0.0038, transition mass 0.2781 ± 0.0038, single-link fraction 0.8717 ± 0.0027, anti-phase-rich occupancy 0.4447 ± 0.0020, and Stopo 0.3038 ± 0.0007. These values place alt3 between the conservative and high-transition references. Importantly, alt3 should be interpreted as an informative transfer test, not as a final optimized biological kernel: `control_base` remains the strongest current hardware anchor for anti-phase-rich occupancy.

### 2.5 Validation-panel correlations identify hardware-aware operational tuning

The validation panel compared `control_base, regular_control, edge_candidate_alt3`, and `overmixed_control` within one 4096-shot run. Statewise exact-vs-IBM ordering was very strong for anti-phase-rich occupancy and Stopo in all four candidates (Figure 5). Stay and Hamming-1 ordering carried the main hardware sensitivity. For `control_base`, validation-panel Pearson *r* was 0.932 for stay and 0.939 for Hamming-1 probability. For `edge_candidate_alt3`, those values were 0.756 and 0.761, respectively. Thus alt3 preserves much of the statewise readout grammar while expressing a more hardware-sensitive transition structure than the baseline.

**Figure 5.**
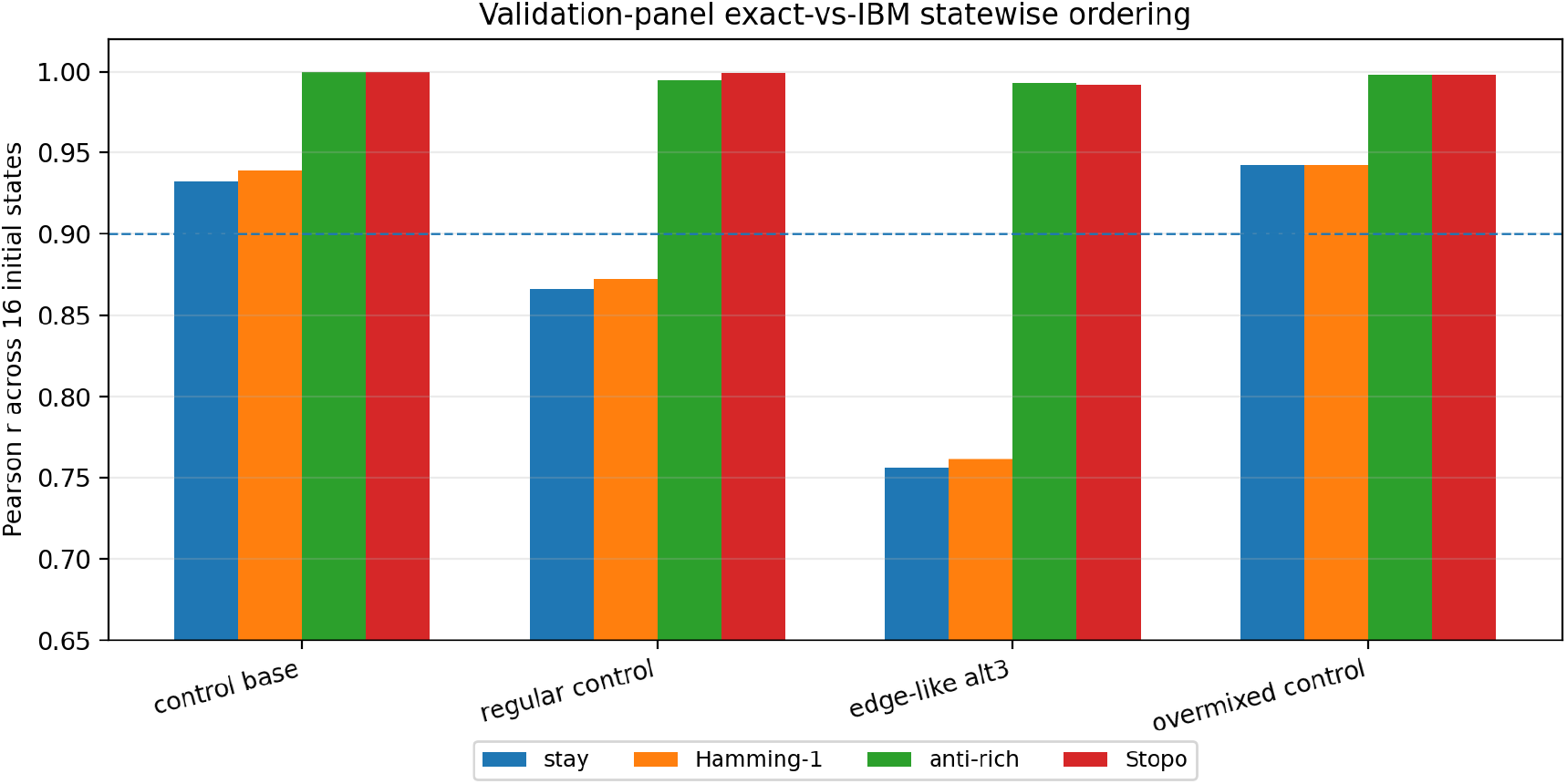
Statewise exact-vs-IBM correlations in the 4096-shot validation panel. Bars show Pearson correlation across the 16 initial topology states for four representative observables. The baseline preserves statewise transition ordering most strongly. The edge-like alt3 candidate preserves high anti-phase-rich and Stopo ordering and marks a more hardware-sensitive transition regime, supporting hardware-aware operational tuning.

This distinction is central to the hardware-aware interpretation. The edge-like candidate is an informative hardware-transfer test: anti-phase-rich and Stopo ordering remain high, transition mass increases, and the aggregate anti-phase-rich readout sits below the locked baseline. The practical design rule is therefore clear. Candidate HSO kernels should be filtered by hardware-aware operational criteria that include statewise ordering, transition mass, one-link fraction, and biology-linked readouts, with exact simulation serving as the first screening layer.

### 2.6 Physiology-linked observables turn hardware execution into an HSO testbed

The hardware observables are tied to specific physiological questions (Table 4). Weighted stay reports how strongly a local coordination pattern persists. Hamming-1 probability reports whether a transition occurs through the smallest possible topological step, a one-pair change along the five-sarcomere chain. Anti-phase-rich occupancy reports whether the local segment occupies states with multiple anti-phase neighbour relations, a feature enriched in the biological HSO topology. The transition edge proxy reports where the minority mismatch pocket lies along the chain. Stopo reports the segment-level synchrony implied by the four neighbouring-pair relations under the equal-amplitude approximation. In this form, the quantum readout is an HSO state-space assay rather than a generic circuit-fidelity score.

**Table 3.** Validation-panel exact-vs-IBM statewise correlations.

| Candidate | Stay $r$ | Hamming-1 $r$ | Anti-phase-rich $r$ | Stopo $r$ |
| --- | --- | --- | --- | --- |
| control_base | 0.932 | 0.939 | 1.000 | 1.000 |
| regular_control | 0.866 | 0.872 | 0.995 | 0.999 |
| edge_candidate_alt3 | 0.756 | 0.761 | 0.993 | 0.992 |
| overmixed_control | 0.943 | 0.943 | 0.998 | 0.998 |

**Table 4.** Physiology-linked observable map. Each quantum observable was chosen to answer a concrete HSO coordination question.

| Observable | Physiological question | Message from the hardware panels |
| --- | --- | --- |
| Weighted stay | Does a local sarcomere phase pattern persist across the short fast-timescale kernel? | The locked baseline and regular reference define a stable conservative region. |
| Single-link fraction | Do transitions proceed by the smallest possible one-pair update? | HSO-like structure is strongest when transition capacity preserves one-link dominance. |
| Anti-phase-rich occupancy | Does the segment occupy local states enriched in multiple anti-phase neighbour relations? | `control_base` gives the strongest current hardware anchor; overmixing lowers this readout. |
| Transition edge proxy | Is the minority mismatch pocket biased toward edge or internal positions? | The model can read mismatch-placement information rather than only global state occupancy. |
| Stopo | What segment-synchrony level is implied by the local topology? | Statewise Stopo ordering remains highly preserved across candidate regimes. |

The biology-side anti-phase-rich target from the five-sarcomere HSO reanalysis is 0.509. The locked exact surrogate gives 0.4513, the 512-shot hardware mean gives 0.4519, and the 4096-shot `control_base` mean gives 0.4517 ± 0.0019. This close exact-to-hardware agreement shows that the baseline readout is hardware-stable. The remaining distance to the biological value is also constrained by the representation: under the fixed empirical prior, the closed-unitary/doubly stochastic upper bound for anti-phase-rich occupancy is 0.4642. Thus the gap to 0.509 is informative model headroom, not evidence that the hardware readout has failed. It marks where additional HSO structure should enter, such as route-aware kernels, condition-aware mixtures, open-system weighting, or slow-fast coupling.

The candidate panels give a concrete design rule. Conservative kernels preserve structure but remain close to a low-transition regime. High-transition kernels add movement but reduce single-link and anti-phase-rich readouts. Edge-like kernels occupy intermediate regions and reveal how exact-screened behaviour transfers to the device. Thus the physiological target is not maximal mixing. It is balanced local reconfiguration: enough transition capacity to represent fast HSO reshaping, enough structure preservation to retain the one-link, anti-phase, and synchrony grammar of the biological topology, and enough hardware stability to keep the statewise readouts interpretable.

## 3 Discussion

This study advances a specific interface question between biophysics and quantum computing: can a local sarcomere coordination grammar, extracted from living-cell HSO data, be executed and interrogated on real quantum hardware while retaining physiological meaning? The experiments support that answer. A five-sarcomere HSO segment becomes a four-qubit local-topology state space; the locked baseline runs repeatably on IBM hardware; and the readouts preserve a direct connection to local HSO reconfiguration, anti-phase enrichment, mismatch placement, and segment synchrony.

The central physiological message is that HSO local coordination is structured rephasing. The relevant behaviour is not random local disorder and not simple global synchrony. It is a regime in which neighbouring relations can change while the biologically meaningful grammar remains local and constrained. This matches the earlier HSO topology in which transitions were dominated by Hamming-1 moves and anti-phase-rich configurations increased during HSOs (Shintani, 2026a). The quantum-hardware panels reinforce this interpretation by showing that excessive transition capacity moves the surrogate away from the HSO-like readout pattern. In other words, the local HSO surrogate is most informative when it preserves structured one-link reconfiguration rather than when it maximizes mixing.

The computational message is equally important. Exact simulation is useful as a first screening layer, but the hardware device supplies an additional selection layer. The edge-like candidate `edge_candidate_alt3` preserved high statewise ordering for anti-phase-rich occupancy and Stopo, yet its aggregate anti-phase-rich readout stayed below the locked baseline. This behaviour is informative rather than disappointing: it shows which parts of the HSO grammar transfer robustly to the device and which parts require hardware-aware tuning. The next model generation can therefore be selected by a physiological score on hardware, not only by exact-simulation performance.

The fixed-prior bound sharpens this interpretation. A closed four-qubit unitary maps the empirical initial-state prior through a doubly stochastic transition matrix, so the anti-phase-rich occupancy reachable under the same prior has a finite ceiling. The computed bound of 0.4642 is below the biological HSO reference of 0.509. Thus the residual gap between the present hardware-stable baseline and the biological target should not be framed as a failure of the IBM readout. It is a representation-level clue: future kernels need biological ingredients that are absent from a closed, static, four-qubit local kernel, such as route-aware history, condition-aware mixtures, open-system weighting, or coupling to the slower beat-scale envelope.

This reframing addresses the main risk for small quantum-hardware studies. A four-qubit circuit by itself would be only a hardware demonstration. Here, the four qubits are the experimentally defined local variables of a living-cell state space, and the observables answer biological questions. The hardware panels ask how the HSO grammar behaves when the local kernel is made conservative, intermediate, or highly transitional. That makes the quantum framework an active assay of physiological state-space structure.

The broader biological interpretation is compatible with sarcomere mechanics. In vivo and cell-level measurements show that sarcomere synchrony and asynchrony influence cardiac performance and persist under living conditions. Oscillatory muscle systems show neighbour-to-neighbour coordination, yielding, and wave-like succession (Yasuda *et al*., 1996; Linke *et al*., 1993; Shimamoto *et al*., 2009; Stehle, 2017; Kagemoto *et al*., 2015). The present quantum-hardware surrogate places the HSO segment into this literature by defining a minimal executable unit of local rephasing. It suggests that fast HSO motion can be understood as a local boundary-shifting process that accommodates phase offsets while preserving beat-scale organization.

The study has a precise scope. It uses one IBM backend, one four-qubit layout, a short Trotter kernel, descriptive repeat statistics, and a local five-sarcomere encoding. It does not claim quantum advantage, a molecular Hamiltonian for sarcomeres, or a complete cell model. Its contribution is a hardware-executed physiological state-space assay. The next advances are concrete: add route-aware mixtures that distinguish compact and extended mismatch motion, add condition-aware kernels tied to beat phase or local amplitude, test cross-backend and device-adaptive layouts, and connect the same observable grammar to simultaneous Ca^2+^, force, or titin-related measurements.

## 4 Conclusion

This study establishes a repeatable four-qubit quantum-hardware surrogate for the fast local topology of HSOs and uses candidate-panel execution to extract a physiological design rule. A useful HSO surrogate must preserve local one-link structure while allowing controlled reconfiguration; simply increasing transition mass is not enough. The locked `control_base` lane is the current hardware anchor, `overmixed_control` defines the high-transition boundary, and edge-like candidates define the hardware-aware tuning path. The result is a concrete bridge from living-cell sarcomere coordination to quantum-hardware state-space testing.

## 5 Materials and methods

### 5.1 Study design and relation to previous biological data

The present work is a quantum-hardware surrogate study anchored to previously reported HSO biology. No new animal or cell-imaging experiments were performed for the quantum-hardware extension. The biology-side definitions of the 16-state local topology, observable targets, and five-sarcomere interpretation were taken from earlier HSO studies cited in the Introduction (Shintani *et al*., 2020; Shintani, 2022, 2026a,b). The new empirical component is the IBM quantum-hardware execution of the locked baseline and candidate panels. All statistics are descriptive and are intended to assess repeatability, statewise transfer, and candidate behaviour within this surrogate family.

### 5.2 Topology reduction and state representation

The observed five-sarcomere segment was reduced to the four neighbouring-pair relations (1–2), (2–3), (3–4), and (4–5). Each relation was encoded as a binary variable, with 1 for in-phase and 0 for anti-phase. The resulting state space consisted of 16 computational-basis states. Given a 4-bit state *s* = (*s*_1_, *s*_2_, *s*_3_, *s*_4_), a five-site sign pattern was reconstructed recursively up to global inversion. Starting from *p*_1_ = +1, we set *p*_*i*+1_ = *p*_*i*_ if *s*_*i*_ = 1 and *p*_*i*+1_ = *−p*_*i*_ if *s*_*i*_ = 0. The topology-derived synchrony proxy was 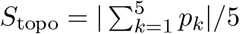. This gauge-fixed reconstruction is sufficient for absolute observables such as Stopo and for edge-versus-centre mismatch proxies that depend only on relative sign structure.

### 5.3 Locked kernel and candidate extension family

The locked static control surrogate was

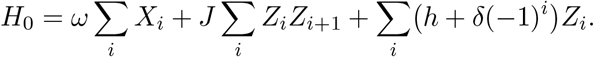

For `control_base`, *ω* = 0.22, *J* = *−*0.8, *h* = 1.0, *δ* = *−*0.2, and total evolution time = 1.0. Candidate extensions added edge-centre, side, and bond-asymmetry terms as defined in the Results. The candidate parameters used in the main validation panel are listed in Appendix Table 7. Circuits implemented a two-step Trotterization of the corresponding kernel.

**Table 5.** Established result and next model layer. The study establishes a hardware-executed HSO state-space assay and identifies specific next biological extensions.

| Component | Established here | Next model layer enabled |
| --- | --- | --- |
| Biological encoding | Five sarcomeres are represented by four neighbouring-pair relations and 16 local topology states. | Longer segments and route-aware mismatch placement. |
| Hardware execution | The 16-state observable pack is executed repeatably on IBM hardware. | Cross-backend validation and device-adaptive layout choice. |
| Physiology-linked readout | Stay, single-link, anti-phase-rich, edge proxy, and Stopo remain interpretable on hardware. | Condition-aware kernels tied to beat phase, $\text{Ca}^{2+}$ context, or local mechanics. |
| Candidate-panel logic | Conservative, edge-like, and high-transition regimes are separated by real-device observables. | Hardware-aware operational tuning of next HSO surrogates. |

**Table 6.** 512-shot per-repeat values for control_base.

| Repeat | Weighted stay | Single-link fraction | Anti-phase-rich occupancy | Transition edge proxy | Stopo |
| --- | --- | --- | --- | --- | --- |
| Repeat 1 | 0.8483 | 0.9095 | 0.4532 | 0.5448 | 0.2938 |
| Repeat 2 | 0.8553 | 0.8973 | 0.4493 | 0.5345 | 0.2955 |
| Repeat 3 | 0.8483 | 0.9158 | 0.4531 | 0.5379 | 0.2945 |

**Table 7.** Main validation candidate parameters.

| Candidate | $\omega$ | $J$ | $h$ | $\delta$ | Time | $\lambda_{ec}$ | $\lambda_s$ | $J_{ec}$ |
| --- | --- | --- | --- | --- | --- | --- | --- | --- |
| `control_base` | 0.2200 | -0.8000 | 1.0000 | -0.2000 | 1.0000 | 0.0000 | 0.0000 | 0.0000 |
| `regular_control` | 0.1588 | -1.3993 | 0.5435 | -0.0742 | 1.9047 | -1.1357 | -0.0891 | 0.5784 |
| `edge_candidate_alt3` | 0.1989 | -1.5229 | 1.3718 | 0.3947 | 2.1728 | -0.3185 | 0.2646 | -0.9213 |
| `overmixed_control` | 0.6715 | -0.7404 | 0.8238 | -0.0985 | 0.5691 | -0.6628 | 0.2971 | 0.3221 |

**Table 8.** Three 4096-shot IBM candidate-panel runs.

| Run | Job ID | Candidate | Weighted stay | Single-link | Anti-rich | Transition mass | Stopo |
| --- | --- | --- | --- | --- | --- | --- | --- |
| run1 | d7ldqqq4lglc7380gbeg | control_base | 0.8387 | 0.9183 | 0.4527 | 0.1613 | 0.2952 |
| run1 | d7ldqqq4lglc7380gbeg | regular_control | 0.8430 | 0.9102 | 0.4482 | 0.1570 | 0.2964 |
| run1 | d7ldqqq4lglc7380gbeg | edge_candidate | 0.6923 | 0.8531 | 0.4383 | 0.3077 | 0.3019 |
| run1 | d7ldqqq4lglc7380gbeg | overmixed_control | 0.6520 | 0.8468 | 0.4336 | 0.3480 | 0.3061 |
| run2 | d87u4n0p0eas73dmmd1g | control_base | 0.8386 | 0.9133 | 0.4495 | 0.1614 | 0.2956 |
| run2 | d87u4n0p0eas73dmmd1g | edge_candidate_alt2 | 0.7068 | 0.8793 | 0.4459 | 0.2932 | 0.3040 |
| run2 | d87u4n0p0eas73dmmd1g | edge_candidate_alt3 | 0.7192 | 0.8698 | 0.4432 | 0.2808 | 0.3043 |
| run2 | d87u4n0p0eas73dmmd1g | overmixed_control | 0.5941 | 0.8087 | 0.4264 | 0.4059 | 0.3104 |
| run3 | d87uengp0eas73dmmp2g | control_base | 0.8414 | 0.9085 | 0.4529 | 0.1586 | 0.2955 |
| run3 | d87uengp0eas73dmmp2g | regular_control | 0.8365 | 0.9044 | 0.4462 | 0.1635 | 0.2975 |
| run3 | d87uengp0eas73dmmp2g | edge_candidate_alt3 | 0.7246 | 0.8736 | 0.4461 | 0.2754 | 0.3033 |
| run3 | d87uengp0eas73dmmp2g | overmixed_control | 0.6027 | 0.8188 | 0.4308 | 0.3973 | 0.3079 |

The exact reference for each candidate was obtained from the same four-qubit Hamiltonian and observable pack for all 16 basis-state inputs. Exact-vs-hardware comparisons therefore test transfer to the intended candidate surrogate within the same model family. Candidate labels such as “regular”, “edge-like”, and “overmixed” are operational labels based on the observable regime, not claims of distinct biological mechanisms.

### 5.4 Observable pack and summary metrics

For each initial topology state, diagonal observables were evaluated for stay probability, Hamming-1 transition mass, Hamming *≥* 2 transition mass, unconditional next-state Stopo, anti-phase count, anti-phase-rich occupancy, edge proxy, side proxy, and transition-conditioned numerators in which the stay state was masked out.

Aggregate metrics were weighted by the empirical prior distribution over initial topology states used in the five-sarcomere HSO reanalysis. Weighted stay is the prior-weighted expectation of the stay projector. Transition mass is 1 *−* weighted stay. The reported single-link fraction is weighted Hamming-1 probability divided by transition mass. The reported transition edge proxy is the prior-weighted transition-conditioned edge-proxy numerator divided by transition mass. To contextualize the biological anti-phase-rich target, I also computed a fixed-prior doubly stochastic upper bound for a closed four-qubit transition matrix; the resulting ceiling was 0.464151.

### 5.5 IBM hardware protocol

All hardware panels reported here used `ibm_pittsburgh` physical qubits [154, 153, 152, 151], EstimatorV2, optimization level 1, and resilience level 1. The 512-shot repeatability panel used 512 shots and repeated `control_base` three times. The candidate panels used 4096 shots. Three 4096-shot panels contained `control_base` and `overmixed_control;` two contained `regular_control;` two contained `edge_candidate_alt3`. Additional exact-selected screening candidates were retained in the analysis records as transfer tests.

For each candidate and each of the 16 initial states, one circuit was prepared and evaluated against the full diagonal observable pack with EstimatorV2. Submission and recovery were separated operationally so that queued IBM Runtime jobs could be recovered later by job ID and matched to candidate metadata.

### 5.6 Summary statistics

Hardware repeat summaries are reported as mean ± sample standard deviation across panels containing the candidate. Because the candidate panels contain small repeat counts (*n* = 2 or *n* = 3 for the repeated candidates), these values are descriptive repeatability summaries rather than inferential confidence intervals. Statewise exact-vs-hardware agreement was computed by comparing IBM and exact values across the 16 initial states using Pearson correlation and mean absolute error. The statistical goal was hardware repeatability, candidate validation, and closeness assessment for a surrogate model family.

## A Per-repeat and candidate-parameter details

### Ethics

This hardware study uses biological source measurements reported in earlier cardiomyocyte studies, for which the original approvals and experimental procedures are described in the earlier publications (Shintani *et al*., 2020; Shintani, 2026a).

## Data and code availability

The reanalyzed dataset, the figure-level summary tables used for the present analyses, and the custom analysis code used in this study are available from the corresponding author upon reasonable request.

## Authors’ contributions

S.A.S.: conceptualization, formal analysis, investigation, methodology, software, visualization, writing–original draft, and writing–review and editing.

## Acknowledgements

Part of this work was conducted as Q0006 (Q-1), a proposal selected in the screening stage of the NEDO Prize Program / Development of Social Issue Solutions Using Quantum Computers (NEDO Challenge, Quantum Computing “Solve Social Issues!”), organized by the New Energy and Industrial Technology Development Organization (NEDO). The author gratefully acknowledges the development environment and mentoring opportunities provided through this program.

## Declaration of generative AI and AI-assisted technologies in the writing process

During the preparation of this work, the author used ChatGPT (OpenAI) to improve the readability and language of the manuscript and to assist with code packaging and tabulation. After using this tool, the author reviewed and edited the content as needed and takes full responsibility for the content of the publication.

