## Supplementary Information for "Repeatable quantum-hardware execution of a fast local-topology surrogate for hyperthermal sarcomeric oscillations"

Seine A. Shintani

#### Scope of this supplement

This Supplementary Information accompanies a preprint manuscript. It is intended as a short reader-facing aid, not as a separate peer-reviewed article or a complete data-and-code archive. It provides only the minimum additional context for the supplementary figures and for the availability of the reanalyzed dataset, figure-level summary tables, and custom analysis code.

#### Supplementary Figure S1

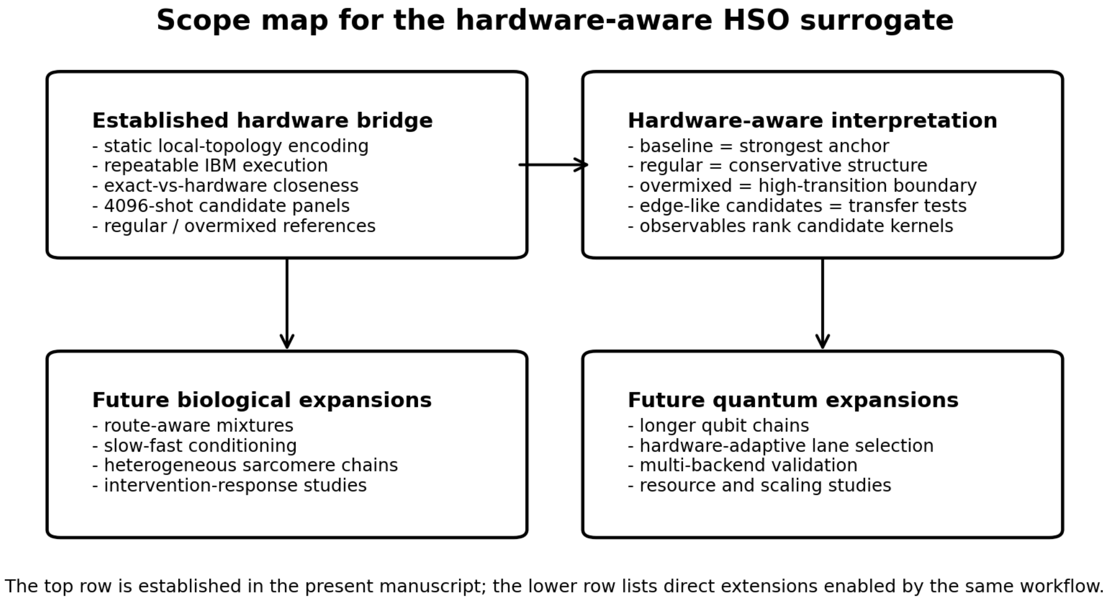


**Supplementary Figure 1: Scope map for the hardware-aware HSO surrogate.** The main manuscript validates the static fast local-topology encoding, repeatable IBM hardware execution, exact-vs-hardware closeness, and candidate-panel references. Future extensions include whole-cell reconstruction studies, resource studies, route-aware kernels, and hardware-adaptive tuning.

### Supplementary Figure S2

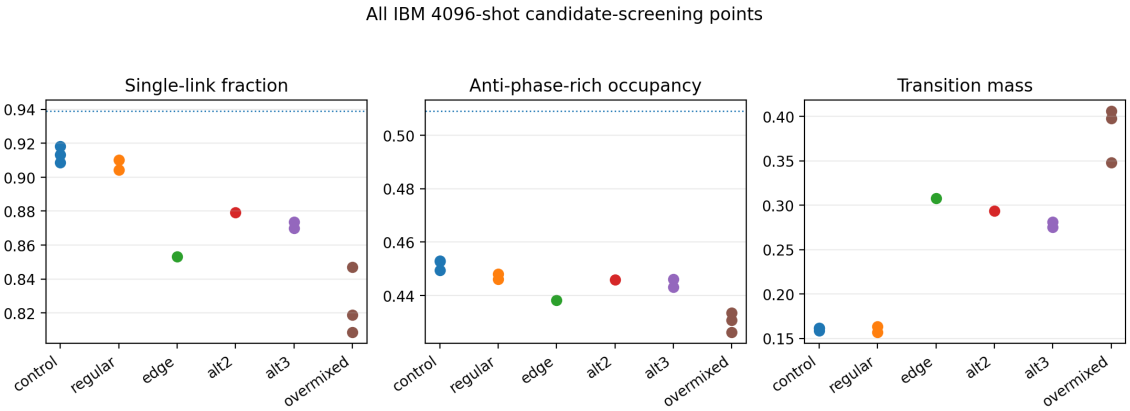


**Supplementary Figure 2: IBM 4096-shot candidate-screening points.** Points show individual IBM runs for candidate labels appearing in the candidate-screening panels. The figure separates the exact-selected initial edge candidate and alt2 from the representative validation-panel candidate alt3 and shows how the panel spans conservative, edge-like, and high-transition regimes.

### Data and code availability

The reanalyzed dataset, the figure-level summary tables used for the present analyses, and the custom analysis code used in this study are available from the corresponding author upon reasonable request.
